# The Role of Retinal Flow in Walking

**DOI:** 10.64898/2026.02.05.704053

**Authors:** Nathaniel V Powell, Daniel Panfili, Youjin Oh, Jonathan S. Matthis, Mary Hayhoe

## Abstract

Both visual direction and optic flow have been implicated in walking towards a goal, in experiments that use prisms to alter visual direction. However, prisms bias retinal flow patterns in ways that may affect balance. This will alter foot placement, and indirectly, a walker’s path. We used a prism manipulation in virtual reality for goal-directed walking, where we manipulated the location of texture-defined flow as well as direction of gaze. The addition of texture straightened paths, but primarily when it provided flow on the ground plane. Texture only on the ground produced straightening comparable to full-scene texture, whereas wall-only texture produced little straightening. This reveals a disproportionate contribution of motion generated on the ground plane to control locomotion, potentially reflecting higher retinal speeds and the prism-induced rotations in the lower visual field. This suggests that, when visual direction is manipulated by a prism, trajectories might result from unexpected motion patterns that activate a balance response, leading to compensatory foot placement. Thus trajectories may be shaped by complex interactions between goal direction, gaze, and local retinal motion patterns.

## INTRODUCTION

Bipedal locomotion is inherently unstable, and one of the requirements of visual guidance of walking is selecting foot placement while maintaining balance. We have relatively little understanding of this visuo-motor control loop. Recent work tracking eye and body movements in the natural world has revealed that the stimulus input to the visual system is dependent not only on the location of gaze but also on the movement of the body. When walking, humans exhibit a sequence of saccades as the body moves forward (Matthis, Muller, Bonnen, & Hayhoe, 2022; Muller et al., 2023, 2024). During the intervening fixations, gaze remains oriented towards a stable location in the world, as illustrated in Figure 1a. This image stabilization is accomplished by the vestibular ocular reflex during which the eyes counter-rotates in their orbit while the body moves forward. The motion projected onto the retina during a step is complex because the head moves up and down and sways laterally during walking. Therefore, the image of the environment on the retina expands and rotates as a consequence of the motion of the head and the reflexive stabilization. This motion occurs along all axes of translation and rotation in a manner that is characteristic of an individual’s gait. This pattern of retinal motion during walking has been measured during natural walking, and can be seen in this Video. This is illustrated at the extremes of the gait cycle in Figure 1b. Because gait patterns are regular, so too, are the retinal motion patterns. Consequently, retinal motion might be used as a dynamic signal to control balance and foot placement during locomotion (Matthis et al., 2022) ultimately influencing the paths that walkers take.

**Figure 1.**
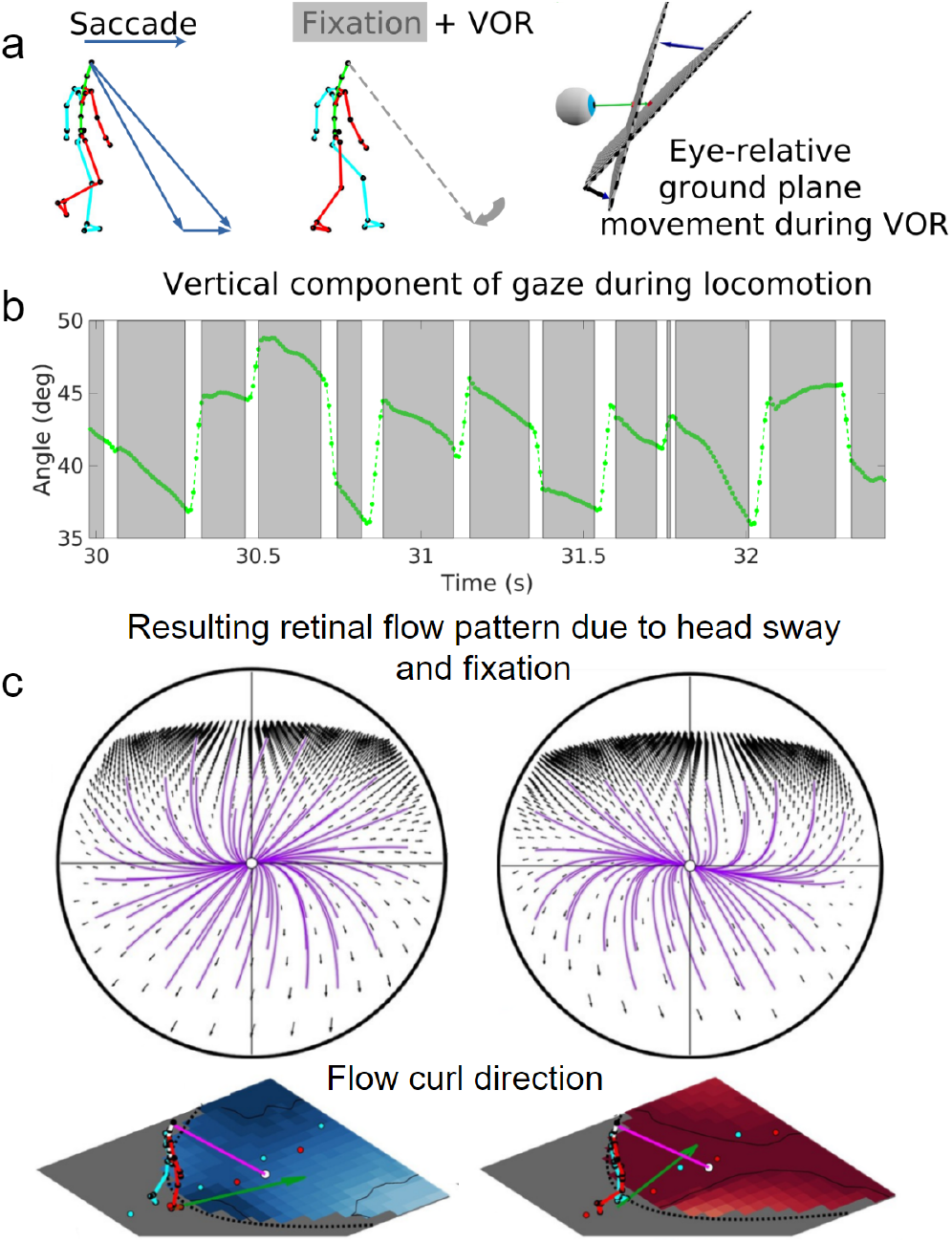
Gaze behavior during a fixation while walking, and the resulting motion signal. (A) Typical fixation behavior while walking, showing the counter-rotation of the eye in the head and how this might lead to retinal motion from the projection of the ground plane onto the retina as the eye counter-rotates in the orbit while the body moves forward. (B) Shows a time trace of the vertical components of gaze angle in the head, showing slow counter-rotation during a fixation. (C) The resulting retinal optic flow pattern (top), which shows two instances (bottom) where the fixation point is to the left or to the right of the current velocity vector of the head (green arrow), respectively. The circular panels show the retinal flow patterns, with the purple lines illustrating the curvature of the flow field and the arrows denoting the direction and magnitude. Fixation to the left of the head’s velocity vector results in a global counter-clockwise (blue) rotation of the retinal flow field and positive retinal curl at the fovea, while fixation to the right of the head’s velocity vector results in clockwise flow (red) and negative curl at the fovea. To see a video of this from real-world walking see https://www.youtube.com/watch?v=Y9NwppteFeE&t=80s

There is considerable evidence to demonstrate that full-field retinal motion exerts a regulatory influence on the placement of the feet. Expansion and contraction of optic flow patterns entrain the gait of subjects walking on treadmills, with the steps phase-locked to the timing of expansion and contraction in the flow patterns (Bardy, Warren, & Kay, 1996, 1999; Warren, Kay, & Yilmaz, 1996). Salinas, Wilken, and Dingwell (2017) also demonstrated regulation of stepping by optic flow, and Reimann, Fettrow, Thompson, and Jeka (2018) showed that optic flow perturbations produce a short-latency response in the ankle musculature. O’Connor and Kuo (2009) showed that optic flow perturbations have a stronger effect when they are in a bio-mechanically unstable direction while standing and walking. It has also been demonstrated that treadmill walkers will rapidly alter their walking speed to match altered optic flow before slowly adjusting back to their bio-mechanically preferred speed (O’Connor & Donelan, 2012). All these studies suggest that retinal flow patterns function as a control variable for stepping. Since walkers have extensive experience with their self-generated retinal motion patterns, it is plausible to suppose that deviations from expected gait-induced retinal motion during locomotion might be used to correct the placement of the feet for greater stability.

In addition to modulating balance, retinal motion is also a very powerful visual cue linked to perception of heading direction. This raises the question of how these two aspects of locomotion work together in natural circumstances. When an observer moves along a linear path, looking in the direction of travel, the retinal motion patterns expand radially, and the Focus-of-Expansion (FOE) of this pattern specifies the current direction the body is moving, or its heading (Gibson, 1950, 1978). It has been clearly demonstrated that observers are able to judge the direction of heading in flow fields that simulate the effect of linear motion, with only 1–2 degrees of error (Warren, 1988). As a result of this and other work, it has been generally accepted that the motion patterns arising from linear or curvilinear translation are used by observers to judge their direction of heading (Burlingham & Heeger, 2020; Chen, Gu, Liu, DeAngelis, & Angelaki, 2016; Dietrich & Wuehr, 2019; Lappe, Bremmer, & van den Berg, 1999; Li, 2025; Li & Cheng, 2011b, 2013; Wall & Smith, 2008). Many experiments on the use of the retinal motion in heading do not involve actively generated body movement, and simulate linear or curvilinear motion, as in steering a vehicle. In this case, there is considerable evidence that the geometry of the flow patterns can be used for perceptual judgments of heading and steering along a linear or curved path (Fajen, 2021; Li & Cheng, 2011a, 2013; Warren & Fajen, 2004). For example, Banks, Ehrlich, Backus, and Crowell (1996) showed that heading can be estimated during eye rotation (while translated along linear paths), but only if the eye movements are real, and generated by the observer. When walking, however, the momentary heading direction varies greatly because of the gait-induced oscillations of the head (Matthis et al., 2022). The presence of gait oscillations has long been recognized (Cutting, Springer, Braren, & Johnson, 1992; Lappe et al., 1999; Lappe & Hoffmann, 2000), but has typically been considered a minor factor (Nakamura, Palmisano, & Kim, 2016; Palmisano, Gillam, & Blackburn, 2000; van den Berg & Beintema, 2000). However, the sensitivity of postural adjustments to optic flow patterns, described above, raises the possibility that the effect of optic flow during natural locomotion may be more complex than usually thought. Others have also pointed out the complexity of the visual signal and argued for its potential role in guiding locomotion (Cutting et al., 1992; Regan & Beverley, 1982).

When walking towards a goal in the natural world, it may not be necessary to use retinal motion or the FOE for heading towards a distant goal. Walkers can use visual direction, that is, the position of a goal relative to the body. Use of visual direction was supported by Rushton, Harris, Lloyd, and Wann (1998), who showed that subjects walking to a goal took a curved path when wearing prism goggles, consistent with the use of visual direction. However, Warren, Kay, Zosh, Duchon, and Sahuc (2001) conducted a similar study in virtual reality, pitting visual direction against the FOE specified by optic flow. This study is particularly important because it involved natural walking, so head movements should destabilize the FOE and add complexity to the flow patterns. In that experiment, visual direction was manipulated by introducing a rotation of the virtual environment relative to the subject while they walked towards a goal. This simulates the effect a prism has on the visual image. The manipulation does not affect the location of the FOE in the absence of gait-induced head movements, and leads to curved paths when walkers use visual direction to control heading. If subjects use the FOE, and head movements are irrelevant, they should walk straight to the goal, whereas if they use visual direction they would follow a characteristic curved path specified by the magnitude of the manipulation. Warren et al. (2001) observed intermediate paths that were straighter than the characteristic curved path predicted simply by the prism’s effect on visual direction. They therefore concluded that walkers use both visual direction and the head-centered FOE generated by optic flow to steer towards a goal, with comparable influences from the two cues.

A complication of the prism is that it affects not only the perceived direction of the target but also introduces a lateral bias in the retinal flow patterns. Prisms introduce an angular rotation of the image and will consequently add a leftward or rightward rotational component to the retinal image during forward translation. This is dependent on the location of gaze relative to the direction of travel. As the eyes counter-rotate during a fixation, a pattern of retinal motion is generated as described above, and the rotational components of that motion pattern will be biased by the prism. When the walker steps toward the apparent location of the goal, the angular displacement of the target is constant, but because walker is closer to the target, its metric displacement of apparent position is smaller, as illustrated in Figure 2. The walker modifies their heading direction because the apparent location of the target has changed, and ultimately walks along a curved path as described in (Rushton et al., 1998). However, it is not only the apparent location of the target that moves laterally as a result of the prism, but the image at other locations in the scene also translates by an amount that depends on distance from the walker. This is the same geometry as that described by Wann and Land (2000) when moving along a curved path, where the retinal motion can be described as curvature in the flow field, or curl. For a full description of the geometry of optic flow, including curl, see (Koenderink & Van Doorn, 1976; Longuet-Higgins & Prazdny, 1980; Regan, 1986). If walkers use the retinal motion patterns to control foot placement, the unexpected motion of the image might induce compensatory foot placement strategies in order to maintain balance, similar to what was shown in (Bardy et al., 1996). If the retinal motion patterns are inconsistent with the normal pattern, it seems plausible that foot placement will be affected, and this could ultimately lead to a change in the walker’s path to the goal.

**Figure 2.**
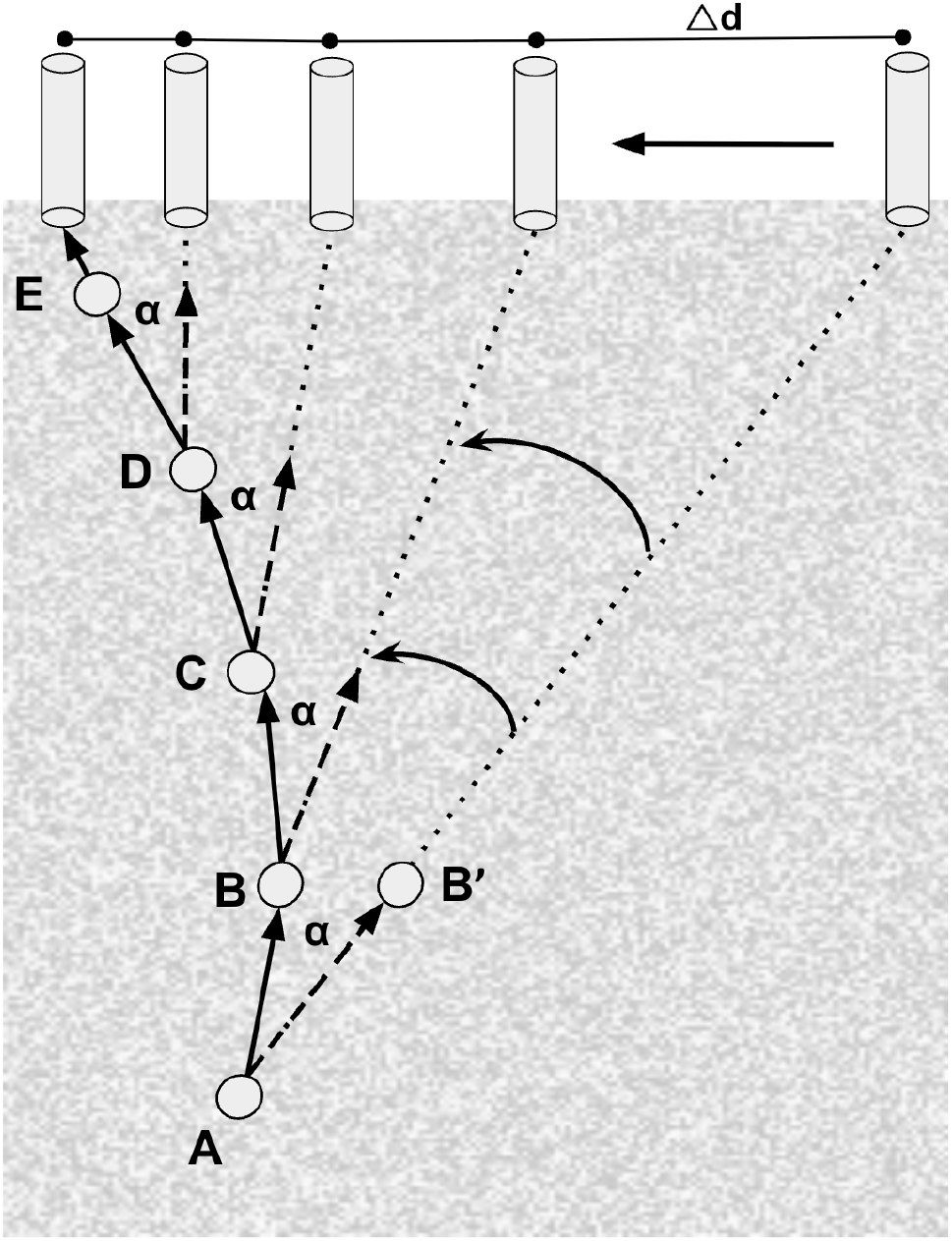
Overview of prism manipulation. The subject starts at A and walks toward the goal (cylinder at the top near position E) whose apparent position is rotated by the prism to the right. As the subject walks forward to B (in apparent direction towards B’), the angle (*α* = 16 degrees) remains constant, but the rate of visual displacement decreases as the walker moves closer to the goal. This process repeats for each step (C-E). This imposes a distance dependent rotation of visual image along the path denoted by the curved arrows, where the texture further away is displaced more than texture closer. This influence diminishes the closer the subject is to the goal.

**Figure 3.**
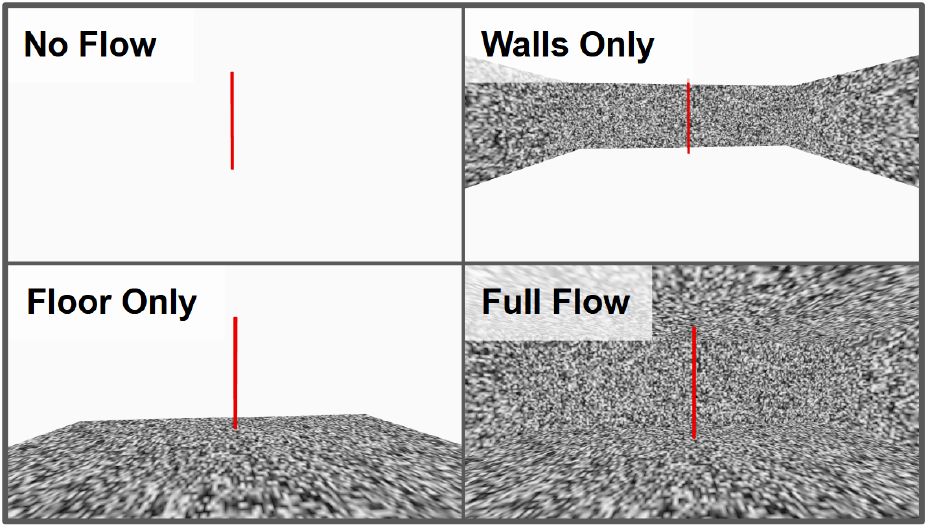
Experimental manipulation of the sources of optic flow. The subject’s goal was to walk to the red cylinder goal object. These are taken directly from the display shown to subjects in VR when facing forward.

Because of the complexity of the retinal motion patterns introduced by the prism manipulation, we reexamined the Warren et al. (2001) experiment in order to investigate the basis of the effect of flow patterns on walking direction. In addition to the distribution of textures across the visual field that were used in that experiment, we added a condition where there was texture only on the walls. We added this condition to explore whether the location of the texture mattered. There is evidence that ground textures are most influential in influencing balance (Fujimoto & Ashida, 2019), and were interested to compare the effect of the floor with the effect of the walls, since both should define the FOE with comparable precision. We also tracked gaze and manipulated where walkers looked in the scene. Walkers typically look primarily at the target in the absence of other instructions. In the current experiment, we added a condition where walkers directed gaze towards the floor. This changes the distribution of motion across the retina and might influence the effect of motion patterns on balance.

## METHODS

### Equipment

The experiment was conducted using a wireless HTC VIVE Pro Eye HMD fitted with a Tobii eye tracker. The eye-tracker cameras had a 120Hz frame-rate, and the HMD screen has a refresh rate of 90Hz. Vive position trackers were used for 6 DOF body tracking for each marker. These trackers were attached on the chest and back, and on the feet. The experiment was developed in Unity (2021.2.10f1) game engine with SteamVR package on a Windows PC (NVIDIA 3090ti GPU and Intel i9 12 core CPU, 64GB 3200MHz memory). The virtual environment consisted of a room (10mx3mx10m) with a green torus as starting zone and a red cylinder as the goal.

Calibration for room orientation and scaling was performed using a custom built calibration routine. Recalibration of the body markers was performed at the start of each trial. An additional calibration step was added for calibrating the eye tracker, provided in the Vive Eye Pro software (eye tracking developed by Tobii). The gaze ray provided by the eye tracker was then projected from the head marker estimate into the three-dimensional environment. The point at which the gaze ray intersected the environment geometry was also recorded in the data collection for further analysis.

### Participants

A total of 14 subjects at the University of Texas at Austin participated in the study. 10 subjects were used for analyses as 4 were excluded due to data recording issues. One subject had a gaze data recording error, and 3 had body tracking failures. All subjects were 20-31 years age, and were balanced equally between male and female. Each individual had normal or corrected-to-normal vision with no known motor impairments.

### Conditions

Subjects were asked to walk to the goal in a virtual room either with or without a prism manipulation, with 16-degree visual offset towards either the right or left. When the prism was active, the visual information was rotated laterally 16 degrees in either the left or right direction relative to the head orientation (along the azimuth). When the walker made a step in the apparent direction of the target, the visual image was updated for the effect of the prism at the current position. Note that as the walker gets closer to the goal, the apparent displacement of the target in meters gets progressively smaller. This is what leads to the curved path characteristic of prisms. As in Warren et al. (2001), an effect of flow was introduced by adding white noise texture to the environment, either to all surfaces, to the walls only, or to the floor only, as shown in Figure 2. In this experiment, the Walls Only condition was added to the set of conditions used by Warren et al. (2001).

At the beginning of each trial, subjects saw a visual cue that informed them to look at either the floor or the goal. The visual cue appeared in the lower visual field 1.5 meters at the beginning of a trial, then disappeared. Subjects were instructed to look towards the ground (even in the absence of texture) roughly 2 steps ahead, which was approximately the distance of the visual cue. Thus gaze location varied as the subject walked towards the goal. The two conditions were labeled Floor Gaze and Goal Gaze. There were a total of 100 trials for each subject, with all conditions given in a random order. The random order was chosen to minimize the prism adaptation effect observed in Bruggeman, Zosh, and Warren (2007). There were 5 no prism trials in each of the environments and 10 trials for each of the other conditions (8 prism direction by environment conditions). There were 10 trials for each environment by gaze condition. Gaze was not manipulated in the No Prism conditions as there is no manipulation to cause deviation from a straight path.

### Data Collection

The data recorded were the condition label, gaze position in world coordinates, tracked body marker positions (feet, arms, torso, and head) and orientations, experiment start time, and frame rate. The data were preprocessed to rotate all position data relative to the start and end positions from the pre-experiment calibration, using a 2D rotation matrix on the x and z axes, orienting the x axis to the path from start to end position and z axis to lateral deviation from the start-end path, keeping the y axis as vertical position (-y = gravity). Trials with leftward prism manipulations were normalized to the rightward trials by inverting the z-axis after applying the calibration rotation.

Pre-processing was performed on the gaze data in order to evaluate whether subjects performed the task appropriately for each condition. A 3D gaze vector was cast out from the recorded head position until it collided with an object in the virtual space. This is marked as the 3D gaze location. Trials with no texture still have collision objects that were rendered invisible to the subjects. Gaze data was marked as being oriented towards the floor if the gaze vector in world coordinates intersected with the floor. The floor texture was rendered invisible in the no flow and wall flow condition, but was still used to determine where the subjects were looking. Because Subjects were walking, eye velocity is non-zero while they fixate, and the eye counter-rotates in the orbit to stabilize against the forward movement. To filter the saccadic eye movements, the angular velocity of the gaze vector in 3D space was checked at each frame. A fixation was detected if the angular velocity was less than 45 *deg/sec* and the angular acceleration less than 15 *deg/sec*^2^ between the frames, and for more than 5 consecutive frames (or 55ms). Note that when subjects are walking, the eye counter-rotated in the orbit to stabilize gaze, so this velocity cut off is higher than for seated subjects.

A data playback tool was used to recover the gaze centered images for each subject’s path trajectory. We used the recorded head position and orientation to place a virtual camera in the virtual space and record the RGB images corresponding to the gaze direction. Using the extracted fixation frames the camera was held stable, oriented towards the initial point of fixation in the virtual space, until a saccade frame, at which point the process started for the next fixation. Within a fixation, average position was estimated across all frames. The camera was then directed to look at this stabilized gaze point, allowing for the removal of noise from the eye tracker. The simulated eye camera was moved forward using the corresponding head tracked data, and thus rotated, as the eye would, while the subjects walked through the virtual space. At each camera position, the RGB image was recorded. The sequence of images for each trial were used to calculate retinal flow (dense optic flow via Deepflow algorithm in OpenCV) which is needed for calculations of curl.

All statistical analyses that compare behavior between conditions were calculated in R using ezANOVA from the ez package. All statistics that required correction for sphericity used Mauchly correction. The between-subject variation was controlled for by treating the subject as a random effect. All multiple comparisons were corrected using Tukey HSD method. Trials in the Floor Gaze condition that contained less than 80 percent of frames with gaze at the floor were excluded from analyses.

## RESULTS

### Path trajectories vary based on flow conditions

For each condition, we calculated the average path trajectory across trials and subjects using the position of the head marker. The last 1 meter was excluded due to noise in position estimates near the boundary of the virtual space. These average paths are shown in Figure 4. As expected, the prism produces curved paths. However, the magnitude of the curvature seems to depend on the location of the texture that produces flow. The Full flow and Floor flow appear to be equally effective in straightening the paths, whereas the Wall flow only straightened the path to some extent when participants look at the goal.

**Figure 4.**
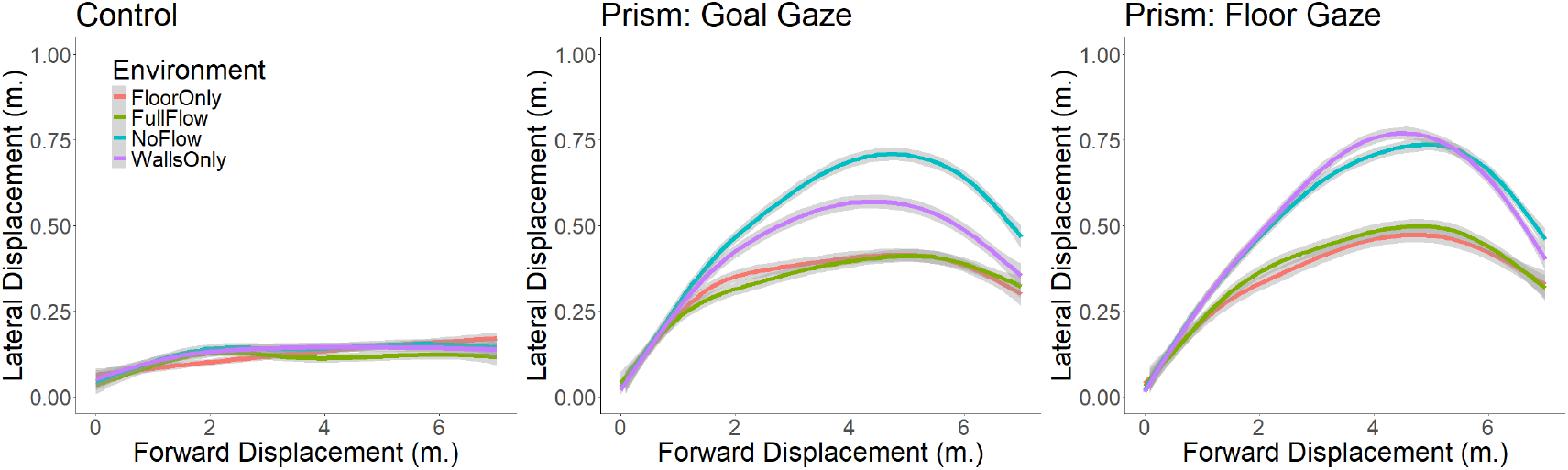
Average path trajectories of the head marker across gaze and flow conditions. The center panel corresponds to when gaze was directed at the goal, and the right panel corresponds to when gaze was directed at the ground. The trajectories in the leftward prism manipulation were flipped over the y-axis so both right and left prism conditions could be averaged together. The gray region corresponds to the 95 percent confidence interval estimated using between subjects variability. These regions are not clearly visible because the confidence intervals are narrow.

To evaluate this in more detail, we calculated the maximum lateral deviation of the paths, defined as the point of maximum deviation left or right of the center line connecting the start position to goal on each trial. The absolute value of this maximum deviation was calculated for each trial, and then averaged within subject and condition. The left and right prism directions were averaged together. The left prism paths were flipped over the lateral axis so that they were in the same space as the right prism path. The control condition deviation is above zero because the unsigned lateral deviation was used. Therefore this reflects variability of the head while walking along a straight path. Similar to Warren’s experiment, we found that paths became straighter as texture (i.e., more flow) was added to the environment. This can be seen in both the average trajectories in Figure 4 and in the average lateral deviations in Figure 5. The presence of texture on the walls only was less effective at straightening trajectories, and does not appear to have any effect when subjects were looking at the floor. This result may stem from the reduction in area of texture visible in the field of view of the helmet when looking down at the floor where there was no texture. The results of a two-way ANOVA show a significant interaction between the flow and gaze conditions (F (6, 132) = 4.541, p *<* 0.001, *η*^2^ = 0.17 [0.06, 1.00]). Post hoc comparisons show no significant differences between the two gaze conditions (t(132) = -0.64, p = 0.80), except for the Walls Only condition (t(132) = 3.375, p = 0.043, d = 1.35 [0.413, 2.289]). No difference was found between the Floor Only and Full Flow conditions. The difference between the No Flow and Walls Only conditions was not significant for either gaze condition, despite the significant effect of gaze in the Walls Only condition, suggesting only a small and variable effect of wall texture.

**Figure 5.**
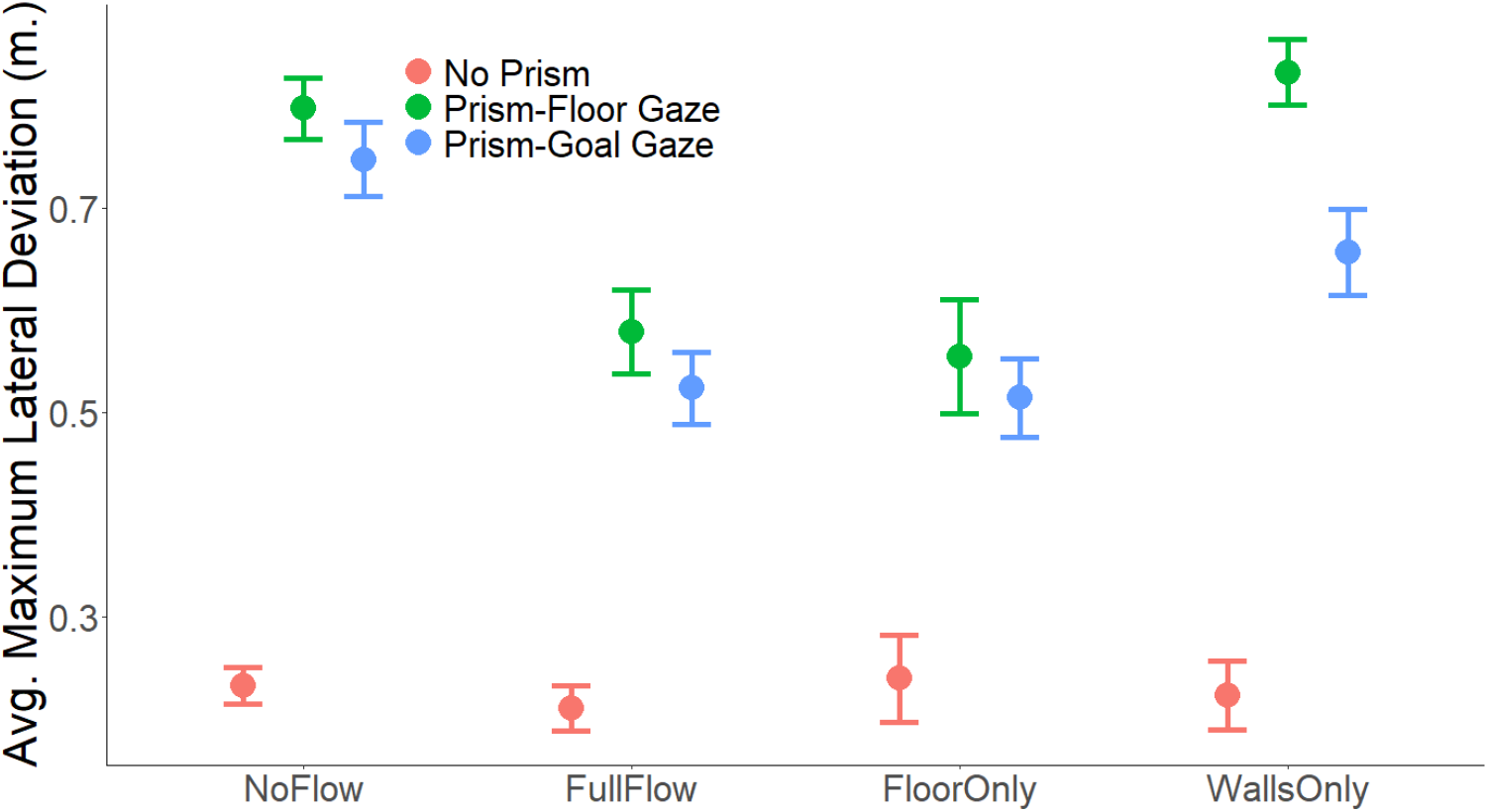
The average maximum lateral deviation (unsigned - left and right are treated the same) of subject head trajectories across conditions. Lateral deviation of the No Prism condition reflects the variability of head sway while walking along a straight path. All error bars are +/-one standard error between subjects.

Straighter paths are evidence of deviations from a pure visual direction strategy. In order to complement the data in Figure 6 and to directly estimate the magnitude of the straightening effect for individual subjects. (While this analysis is similar to the previous one, it allows removal of the between-subject variability.) In this analysis, the prism/no texture condition was treated as a control. For each subject, the average path in each texture environment was calculated. At the halfway point for each average path (4m) the difference between the No Prism and the texture condition was extracted for analysis (See the red dot in the left panel of Figure 6). This point was chosen as paths were most curved at approximately this point on average.

**Figure 6.**
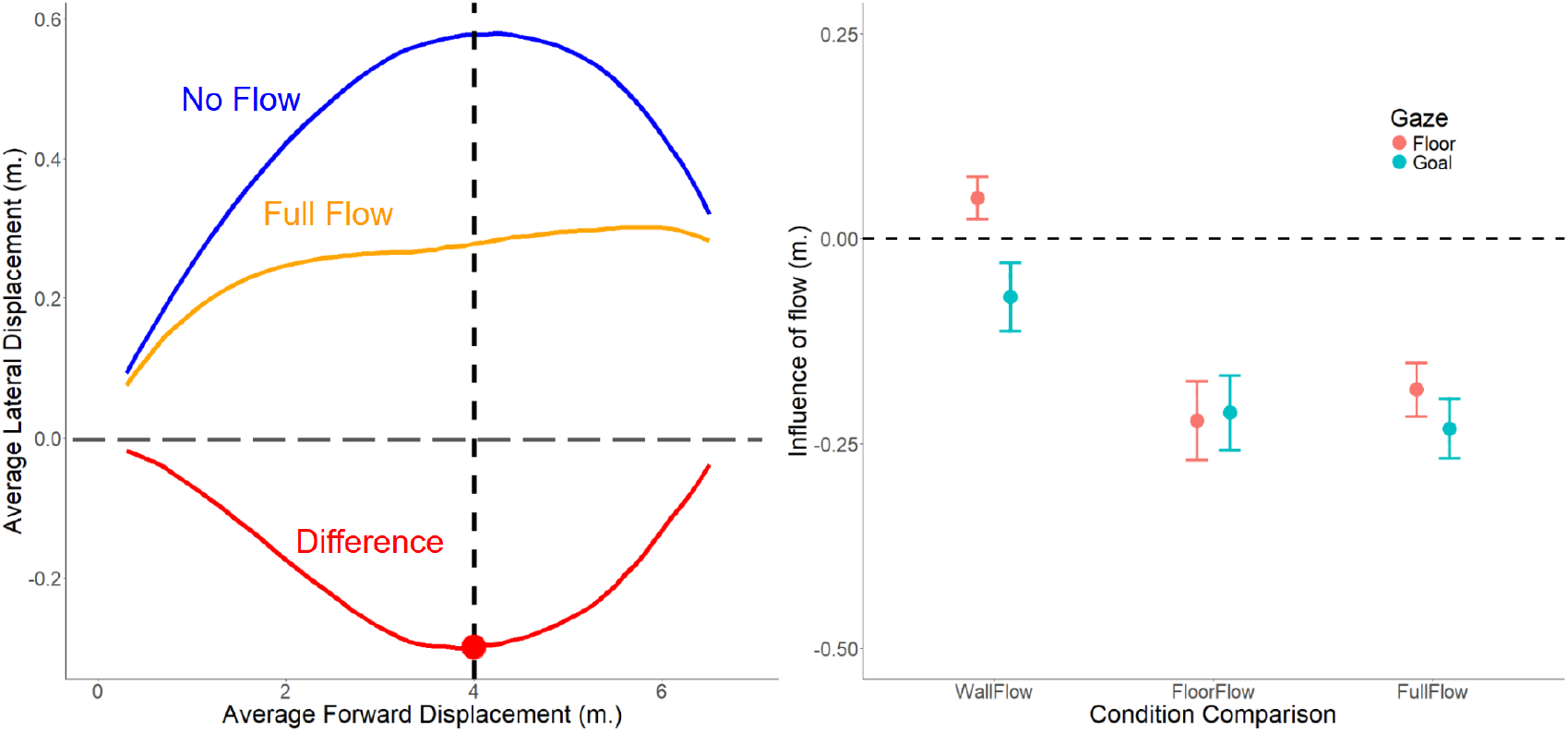
Within subject analysis of the effect of Flow conditions to straightening paths. (Left) An example of how differences between path trajectories were calculated for a subject. The effect of the prism with no texture was treated as a control, and the difference between the control trajectory and the wall flow trajectory is shown by the red line. Negative values indicate a straighter path. 4m was the halfway in each trial. The red dot corresponds to a point taken at 4m. (Right) The average difference in path trajectories in the two gaze conditions. The points refer to the average difference at 4m (red dot) across wall, floor, and full flow conditions for each subject. All error bars are +/-one standard error between subjects.

This comparison, shown in Figure 6, shows that Floor and Full Flow conditions have a substantial effect of straightening the path in both gaze conditions. The results from a one-way ANOVA shows a significant main effect of flow condition (F(2,33) = 13.593, p *<* 0.0001, *η*^2^ = 0.45 [0.23, 1.00]). Post hoc tests reveal that there is no difference between Floor Only and Full Flow conditions (F *<* 1, p = 0.84). A significant difference was found between Walls and the other two texture conditions (F(1,33) = 27.142, p *<* 0.0001, *η*^2^ = 0.45 [0.24, 1.00]). While there is a small deviation from zero in both Walls conditions, they are in opposite directions, and not significantly different from 0 in either gaze condition.

### Head-Foot coordination is biased by the prism

To examine whether the prism influenced the behavior of the feet, we looked at the angle between the head and foot direction. The relative orientation of the forward facing head and each foot was calculated for each frame, and then the average angle was used for analysis. Figure 7 plots this angle relative to the No Prism condition. The figure inset illustrates this angle. This analysis shows that the presence of floor texture changes the angle between the head and feet. Using a linear mixed effects model results in a main effect of the environmental textures (F(3,186.84)=16.5, p*<*0.001, *η*^2^=0.21[0.12,1.00]). In both the Walls Only and No Flow condition, the angle is smaller than both the Floor Only and Full Flow conditions. The results of a post hoc comparison from planned contrasts between the two floor texture containing conditions and the two other conditions show a significant difference (t(188)=7.00, p*<*0.0001, *d*=0.97 [0.698, 1.246]). The sign of the angle indicated that the feet oriented inside of the direction of travel. This shows that the feet are most affected in the conditions where the paths were most affected by retinal motion.

**Figure 7.**
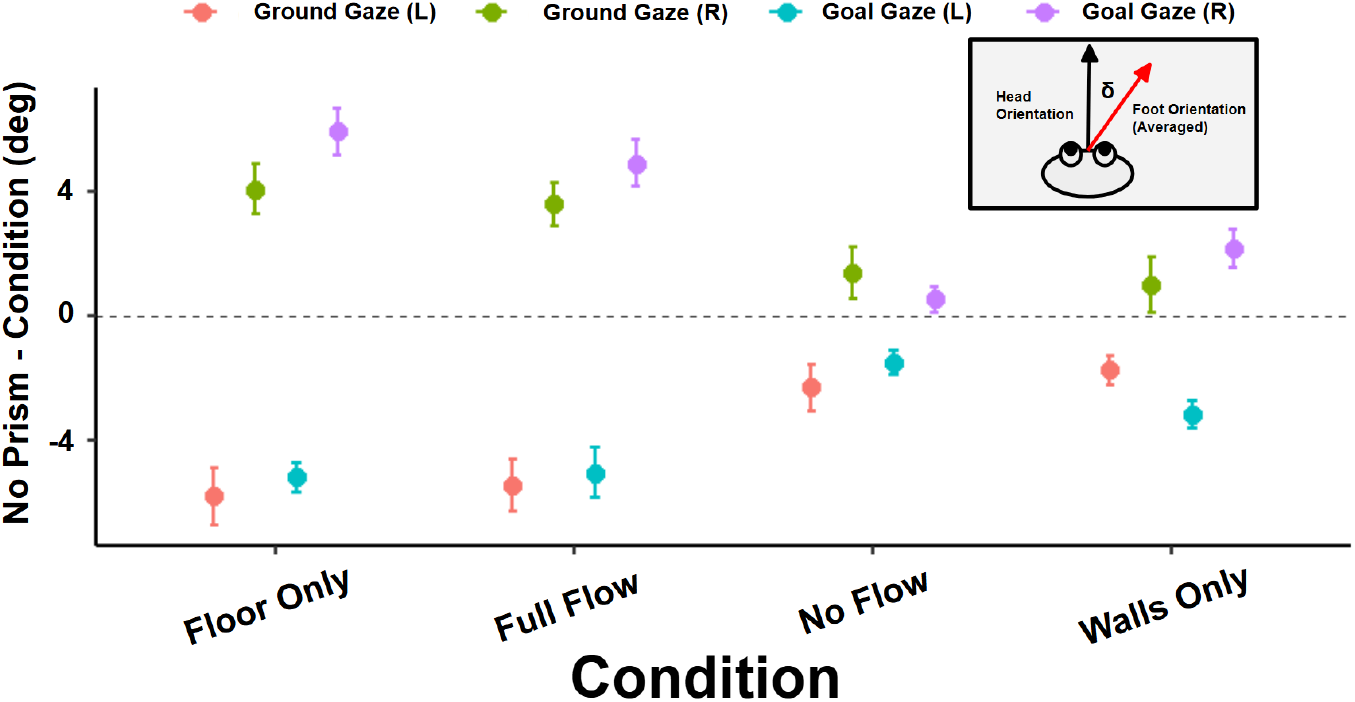
Angle between head and foot orientation across conditions. Each point corresponds to the angular difference between the head and the feet (averaged). The small inset illustrates this angle and denotes it as *δ*. Error bars are +/-1 standard error.

### Retinal Flow Patterns

While our results on the effect of added texture in these conditions are similar to those of Warren et. al (2001), we find, in addition, that the effect of the flow appears to be coming largely from the texture on the floor. Flow from the floor alone was as effective in straightening the paths as flow present on all the surfaces (i.e., Full Flow). There was a small effect of the Walls Only texture when subjects looked at the goal. In the Introduction, we described how the prism might add distance-dependent lateral motion to the image (See Figure 2), and suggested that unusual retinal motion patterns might affect foot placement if retinal flow controls posture and balance. We therefore quantified the average retinal motion component added by the prism manipulation. This is shown in Figure 8. The goal of this calculation was to summarize the retinal flow patterns during the first half of the trial.

**Figure 8.**
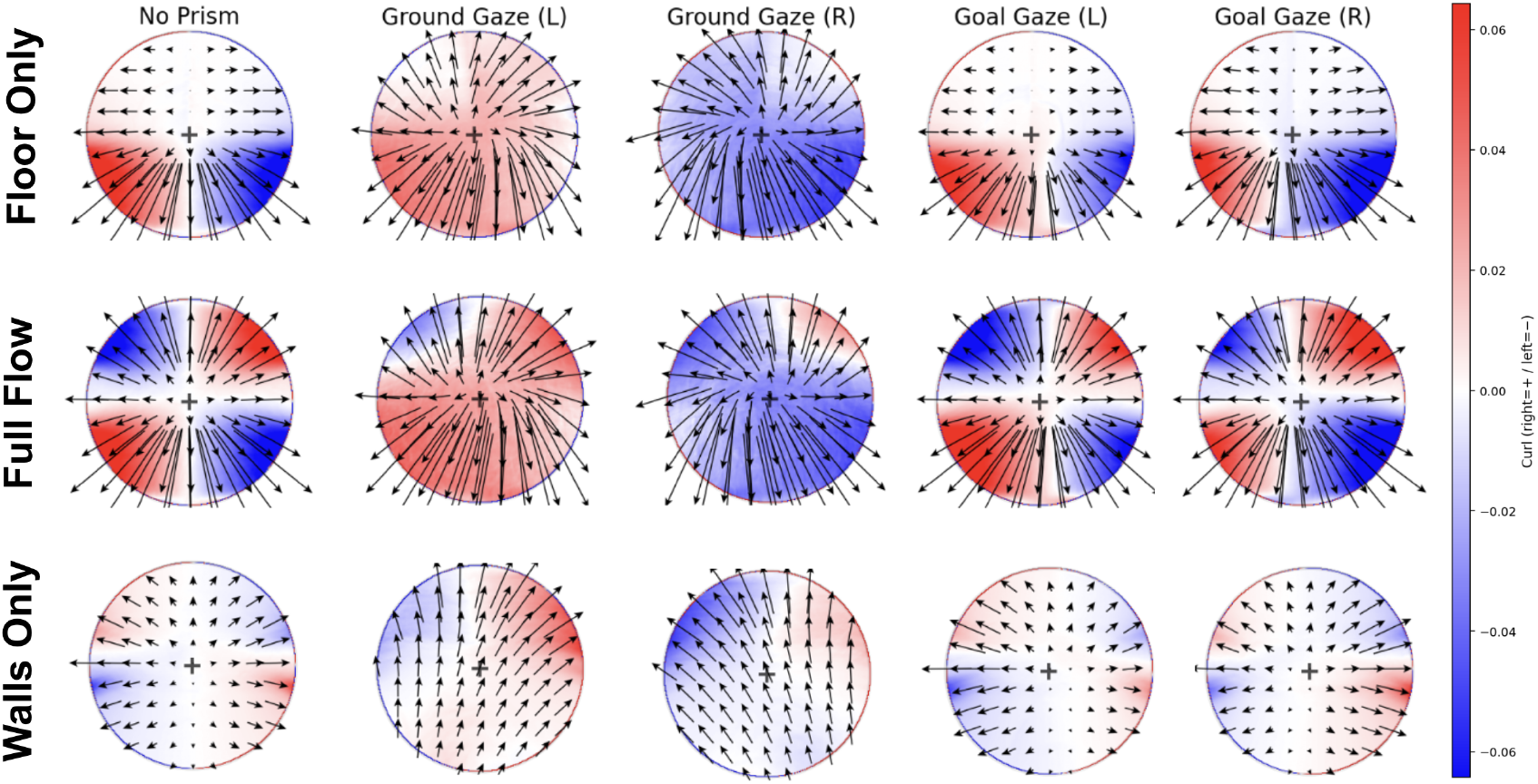
Each circular panel depicts the average curl across an 89 degree window centered on the gaze point. For each condition, the retinal flow was averaged within gaze and texture conditions. For each condition, only the first half of each trial was averaged, where the prism effect is strongest. The black arrows denote the average local velocity. The colors specify whether curl is positive or negative. White regions correspond to areas with zero curl.

Retina-centered flow was calculated by recording the sequences of images in Unity that corresponded to the exact image sequence subjects saw during the experiment. The head camera was rotated to look at the gaze point to simulate the visual stimulus on the retina. At each recorded head position, the gaze centered camera view was recorded. For each frame of the gaze centered video, the RGB image was recorded with vertical field of view 89 degrees and horizontal field of view 120 degrees, and 1080×1920 pixel resolution. Optic flow was calculated from the resulting images using the OpenCV’s Deepflow algorithm, using only the fixation frames defined by angular velocity of the eyes rotation. We then averaged these vectors over the frames in a trial, then over trials and subjects. Only frames corresponding to fixations were used, but the retinal image changes as subjects walk, get closer to the goal, and shift gaze to new locations. Note that the flow patterns here are averaged across the head positions associated with body sway, and simply reflect the bias induced by the prism. The frames from the first 50 percent (first 4m of walking) of each trajectory were used for analysis, since this is where the displacement of the target by the prism is greatest. Figure 8 shows the average retina-centered flow for each of the three texture environments. These patterns are a simple effect of the geometry. In the Figure, the arrows show the direction and magnitude of the local retinal motion vectors. The colors reflect the magnitude of the rotational component, where red is positive curl (clockwise rotation) and blue is negative (counter-clockwise rotation). White indicates zero curl. This figure helps visualize the effect of the prism averaged over body sway given the location of gaze in the trial. Looking at the Full Flow condition, it can be seen that the prism introduces a strong rotational component in the ground gaze condition indicated by the direction of the motion vectors and the location of the region of zero curl. There is a less obvious rotational component when walkers look at the goal, which is most easily seen by the displacement of the zero curl region. Similar effects are seen in the Floor Only condition, but are restricted to the lower visual field when looking at the goal. In the Walls Only condition the region of zero curl is similarly displaced, but the magnitude of the motion vectors is reduced, as there was no texture on either floor or ceiling. In addition, the retinal image shifts within a trial and will vary with subjects and over trials. Additionally, this smears textured and untextured regions. These patterns are similar to what was calculated by Matthis et. al., (2022), shown in the bottom panel of Figure 1.

Figure 8 demonstrates the complex retinal flow patterns in the experiment that are induced by the prism manipulation. In particular, the retinal location where there is zero curl (on average) is biased depending on the direction of the prism. This bias will be added to the gait-induced motion resulting from body sway, and the unexpected motion might signal an error in balance and lead to corrective placement of the feet. We cannot make a direct connection between the flow patterns and the placement of the feet, given that we averaged over a number of steps in a section of the trial, and a response to an unexpected retinal motion pattern is likely to operate on a step-by-step basis. This Figure simply demonstrates the overall bias caused by the prism and gaze location. We do not know how the visual system weights the retinal motion information across the visual field. Full Flow and Floor Flow affect path curvature in a similar manner, regardless of gaze location. Therefore it is interesting to note that the retinal motion patterns in these conditions are different, although the location of zero curl (white regions, where there is approximately zero rotation of the motion vectors) is similarly biased in direction. This result suggests both that the response to the unusual motion patterns is complex, and that motion from the floor appears to be most important. The retinal motion patterns are also consistent with the attenuated effect of the Walls Only condition. When gaze is on the goal, the region of zero curl is displaced as expected by the prism manipulation. The speed of the motion is attenuated, perhaps accounting for the somewhat smaller effect of the prism on path straightening. When subjects look mostly at the floor, the texture from the walls will be in the upper visual field. In this condition, subjects did not appear to straighten their paths, consistent with the point made above that the Floor texture is most effective.

## DISCUSSION

The goal of this experiment was to re-examine the effect of flow in straightening paths in the presence of a prism. Prisms rotate the visual image and bias retinal motion as a walker approaches a fixed goal. This generates a complex pattern of motion on the retina. There is considerable evidence that optic flow patterns influence balance and the placement of the feet during locomotion Lee and Aronson (1974); Lee and Lishman (1975); Mohler, Thompson, Creem-Regehr, Pick Jr, and Warren Jr (2007); Prokop, Schubert, and Berger (1997); Salinas et al. (2017). There is also evidence that the motion patterns walkers normally generate are linked to their gait in a regular manner (Matthis et al., 2022). If walkers learn these regular patterns, it seems likely that the unexpected retinal motion caused by the prism might be interpreted by the walker as a balance problem and lead to changes in foot placement.

The path trajectories we observed were comparable to those found by Warren et al. (2001), in that the addition of surface textures straightened the paths the subjects took. In our experiment we added a condition not in the original Warren experiment, where texture was confined to the walls. Notably, we found the addition of texture alone is not enough to produce a straightening effect. We found that the straightening effect of texture on the floor is comparable to that of texture on all the surfaces. The straightening effect did not appear to be greatly influenced by gaze location in these conditions. However, the effect of texture on the floor was much greater than the effect of texture on the walls. This is consistent with observations of a greater effect of the lower visual field on postural control (Fujimoto & Ashida, 2019). While the origin of this effect is unclear, it might be a consequence of the higher speeds and greater curl of the motion vectors close to the body. In contrast, the presence of flow from the walls alone produced paths only slightly straighter than with no texture, when walkers looked at the goal. This effect was statistically significant in Figure 5 but not in Figure 6, so it is hard to know how much confidence to place in this result, and therefore we treat it here as a small effect. In the condition where walkers looked at the floor, the textures fell primarily in the upper visual field, and only a small area was visible. Consequently, there was no straightening of the path, consistent with the reduction in local motion speed and rotation on the floor. This appears consistent with Zorpala and Lopez-Moliner (2026) whose evidence suggests that biases in the curl signal are responsible for perceptual biases in heading beyond the FOE.

The goal of the current experiment was to explore how motion patterns caused by the prism might influence paths taken in the presence of a prism. Although it is difficult to establish a direct causal link between the paths chosen and the retinal flow resulting from the prism manipulation, the literature suggests that an abnormal flow pattern is likely to influence foot placement on a step-by-step basis. The patterns shown in Figure 8 are averaged over steps and trials, and are affected by any changes in gaze location that occur over the first segment of the trial. To make a closer link it would be necessary to simulate the experiment for each step and at a particular gaze location. In addition, it would be necessary to estimate parameters for the weighting of retinal motion across the visual field, as well as for a walker’s sensitivity to deviations from their normal gait-linked flow. The main conclusion that can be drawn from our calculations of retinal motion is that the prism manipulation introduces biases in the retinal motion patterns that may influence foot placement, and thereby the path chosen.

Thus the effects of the prism in both our experiment and that of Warren et al. (2001) appear to be more complicated than a weighted effect of visual direction and the FOE. The effect of visual direction on walking towards a goal is clear, but the way that the retinal motion patterns affect paths appears to be quite complex. The chosen paths appear to result, at least in part, from a balance response to unusual retinal motion, and may not be closely linked to steering towards a goal. We also showed in Figure 7 that the head and torso was consistently pointed towards the apparent position of the goal, but the feet were more often misaligned with that direction, instead pointed to the inside of the current path. This suggests corrective foot placements and changes in posture. This effect was more prominent in the two conditions that contained the ground plane, suggesting its link to making foot placement decisions.

In a related experiment using a prism, Harris and Carré (2001) reported straighter paths when subjects reduced their viewing height by crawling toward the target. They suggested that this effect might come from changes in available texture information across the visual field (Harris & Carré, 2001). This distinction is useful for interpreting our results. In our VR setup, the field of view was comparatively large and the ground plane was visible, so their explanation, that the effect of the prism is caused by loss of the foreground in their goggles, is unlikely to be a primary driver. However, their result does give a plausible account of how changes in posture/eye-height can alter the retinal flow field and its utility for control. However, it is unlikely that the artifact they report would be present in our experiment. We show that the ground is indeed important, but this is because this is where the influence of the prism on the flow signal is strongest.

It was somewhat surprising that paths were largely not influenced by gaze position except in the walls-only condition. One interpretation is that, under full texture or ground-only texture, the locomotor system can obtain sufficient ground-generated motion even when gaze is directed towards the goal, because the lower visual field still contains robust ground-plane structure and because walkers can keep their body oriented toward the goal even while sampling near-ground information peripherally. By contrast, in the walls-only condition, looking down removes most of the remaining texture from view (due to head pitch), sharply degrading the available motion structure, and so the straightening effect is partially removed. This dependence on visual texture in different parts of the visual field suggests that there is not a uniform global FOE computation for instantaneous heading, but rather a reliance on weighting regions of the visual field differently.

The FOE has historically been treated as the primary cue for guiding direction of locomotion, motivating texture manipulations intended to strengthen or weaken global optic flow. However, a wide set of steering studies suggest that the visual information used for control is often not the FOE per se, but information about the future path that can be sampled from specific regions of the flow field and stabilized by gaze. For example, observers can judge a point on their future path more accurately than an instantaneous heading during curvilinear motion, which implies that path-specifying variables can be more reliable than heading when rotation and curvature are present (Wilkie & Wann, 2006). Extending this, other studies have shown that active look-ahead strategies provide an effective control variable for steering around obstacles, formalizing how the control system can rely on sampled visual information rather than a single global flow feature (Wilkie, Wann, & Allison, 2008). In driving, Lappi (2014) argues that steering around bends is naturally described in terms of information about the upcoming trajectory, with gaze behavior organizing which sources of information dominate moment-to-moment (Lappi, 2014; Lappi et al., 2020). This perspective maps directly onto walking, where the oscillatory head movements due to gait make the retinal signal different from the smoother flow experienced in simulated or vehicle steering, increasing the importance of where in the scene information is available. Thus, visual direction to the goal can serve as a far-point orienting reference, while retinal flow over the ground plane supplies the near-point information needed for rapid stabilization and step-by-step foot placement, consistent with the near–far control logic of the two-point steering model (Salvucci & Gray, 2004). Under this viewpoint, our texture effects follow from the source of usable information, where ground texture does not merely add flow, but provides dense, diagnostic ground-referenced flow lines that specify movement over the walking surface and support moment-to-moment control. This is further supported by Cutting (1992)’s emphasis that near-ground retinal-flow invariants disproportionately support guidance during gait (Cutting et al., 1992). Under prism-induced retinal-flow biases, preserving near-point information provided by the ground appears to yield similar path straightening in the Full Flow and Floor Only conditions. However, in the Walls Only and No Flow conditions, near-point ground information is largely removed. This could be interpreted as pushing walkers towards greater reliance on far-point information, which is associated with less accurate movement control (Salvucci & Gray, 2004).

Gaze while walking is comparable to that in driving, in that near and far looks may serve a different purpose (Lappi & Mole, 2018; Salvucci & Gray, 2004). In natural settings, walkers alternate between making eye movements to the ground and into the distance (Matthis, Yates, & Hayhoe, 2018). On flat surfaces subjects look into the distance roughly 50 percent of the time, and this becomes much less frequent as the terrain becomes more rugged. In rugged terrain, walkers mostly allocate gaze to the ground in front of them to guide foot placement, and only occasionally look in the distance (Matthis et al., 2018). This suggests that using visual direction for steering along a path is not very demanding, and occasional looks are adequate. Thus gaze in the distance and gaze on the ground may serve different purposes, with gaze in the distance being used for controlling walking using visual direction to the goal (i.e., heading) and gaze on the ground used for control of step placement and balance, using retinal flow (Matthis et al., 2022; Muller et al., 2024). Thus the effect of retinal flow patterns on paths in this experiment seem most closely linked to balance, and does not strongly support the use of the FOE in controlling direction toward a goal.

